# Reticulon dependent ER-phagy mediates adaptation to heat stress in *C. elegans*

**DOI:** 10.1101/2024.07.26.605287

**Authors:** Vincent Scarcelli, Claudia Serot, Alexandre Pouget, Céline Largeau, Audrey Sagot, Kenza El-Hachimi, Denis Dupuy, Emmanuel Culetto, Christophe Lefebvre, Renaud Legouis

**Author notes:** Equal participation (Vincent Scarcelli, Claudia Serot).

## Abstract

The selective degradation of ER by autophagy, named ER-phagy, promotes the recovery of ER homeostasis after a stress. Depending on the ER stress, different types of ER-phagy involve various selective autophagy receptors. In this study, we report a macroER-phagy induced by the fragmentation of tubular ER in response to acute heat stress. We identified a novel ER-phagy receptor encoded by the reticulon long isoform RET-1d. RET-1d is mainly expressed in the nervous system and the epidermis and colocalize with the ubiquitin-like autophagy protein LGG-1/GABARAP during heat stress induced autophagy. Two LIR motifs in the long intrinsically disordered region of RET-1d mediate its interaction with LGG-1 protein. The specific depletion of RET-1d isoform resulted in a delay in autophagosome biogenesis and a decrease in the capacity of animals to adapt to heat stress. Our data revealed a RET-1d dependent ER-phagy mechanism that takes place in neurons and epidermis and participates to the adaptation of *C. elegans* to heat stress.

## Introduction

The endoplasmic reticulum (ER), the largest membranous organelle in eukaryotic cells, endorses multiple functions among which the synthesis, folding and quality control of proteins, the biosynthesis of lipids, and the regulation of calcium homeostasis (Schwarz & Blower, 2016). Depending on the presence of ribosomes, ER is qualified of rough (rER) in opposition to smooth ER. The structure of the ER is complex and very dynamics (Nixon-Abell *et al*, 2016), forming a network of dense tubular matrices and tubules which establishes numerous membrane contact sites with other organelles and the plasma membrane (Phillips & Voeltz, 2016). The morphology of the ER, which is tightly linked to its functions and constantly undergoing rearrangements, is regulated by a series of proteins that control the level of curvature, the tubules fission or fusion, and the stabilization of contacts (Obara *et al*, 2023; Shemesh *et al*, 2014). The ratio between sheets and tubules varies depending on the cell type and functions (Westrate *et al*, 2015), for instance axonal ER appears as a smooth continuous tubule that could extend over long distance (Öztürk *et al*, 2020; Kuijpers *et al*, 2023).

The reticulons and reticulon-related proteins family localize mainly at ER membrane and are essential for generating and maintaining the structure of ER-tubules (Voeltz *et al*, 2006; Shibata *et al*, 2010; Westrate *et al*, 2015). The C-terminal reticulon homology domain (RHD) of 150-200 amino acids, which consists of two hydrophobic regions separated by a 66 amino acid hydrophilic loop, adopts a hairpin-like W-topology (Breeze *et al*, 2016), establishing a positive membrane curvature (Zurek *et al*, 2011). Although the C-terminal RHD is well conserved in all species, the N-terminal domains are highly variable in sizes and sequences from a few to over a thousand amino acids. In mammals, the four reticulon genes *rtn1, rtn2, rtn3, rtn4/nogo*, encode for several isoforms through alternative splicing sites and various promoters (Oertle *et al*, 2003). The localization and specificity of these isoforms are yet largely unknown but some of them have been recently involved in autophagy during ER stress (Grumati *et al*, 2017; D’ Eletto *et al*, 2019).

Cellular stresses can reduce the capacity of ER to fold and process proteins, leading to the accumulation of misfolded or unfolded proteins. Altered ER proteostasis has been implicated in the occurrence of a variety of human diseases, including cancer, neurodegeneration, metabolic diseases and chronic inflammation (Hetz *et al*, 2019). ER stress could have a strong impact on proteostasis and Ca^2+^ homeostasis and triggers an adapted response coordinating the Unfolded Protein Response (UPR), the ER-associated protein degradation (ERAD), apoptosis and autophagy. Macroautophagy, commonly referred as autophagy, is a major degradative process that involves the sequestration of damaged organelles, misfolded proteins, and other cellular debris into double-membraned vesicles called autophagosomes. Autophagosomes then fuse with lysosomes, where their contents are degraded and recycled. This process plays a critical role in maintaining cellular homeostasis, responding to nutrient deprivation, and defending against infections and diseases by clearing out harmful or unnecessary cellular components (Klionsky *et al*, 2021). The autophagy is selective when defined intracellular components are specifically sequestered by autophagosomes, in opposition to bulk material, and further named according to the degraded cargoes: mitophagy, lipophagy. Selective autophagy relies on selective autophagy receptors (SAR) which mediate interaction between the cargo and the nascent autophagosome by binding the ubiquitin-like proteins LC3/GABARAP generally through LC3 Interacting Region (LIR) domains (Galluzzi *et al*, 2017).

The selective degradation of ER pieces, which is called ER-phagy or reticulophagy, participates to the regulation of the shape, the contents and the homeostasis of the ER, and is essential during drug-induced ER stresses. Recently, several ER-phagy receptors have been identified, which can be either cytosolic or ER membrane proteins (Gubas & Dikic, 2022; Chen *et al*, 2019). Data in yeast, plant and mammals have shown that reticulons and RHD containing proteins have SAR function for reticulophagy during ER stress (Mochida & Nakatogawa, 2022; Chino & Mizushima, 2023; Reggiori & Molinari, 2022). The mammalian FAM134B and RTN3 proteins participate in remodeling the ER network and the elimination of specific domains by reticulophagy (González *et al*, 2023; Berkane *et al*, 2023; Khaminets *et al*, 2015; Grumati *et al*, 2017). These pioneer works have raised numerous questions on the functions of these SAR concerning the specificity of the isoforms, their interactions with LC3 and GABARAP members, the categories of ER cargoes and the relations with the types of stress. In human, ER-phagy receptors are associated with several neuropathologies such as hereditary sensory neuropathy and Alzheimer disease (Guelly *et al*, 2011; Murphy *et al*, 2012; Zou *et al*, 2018; Kim *et al*, 2023; Foronda *et al*, 2023; Kurth *et al*, 2009). However, the presence of multiple reticulon and RHD proteins as well as several LC3 and GABARAP members complicates the analysis of their specific functions.

In this study, we characterized in the nematode *C. elegans* the unique reticulon protein RET-1 (Oertle *et al*, 2003; Yang & Strittmatter, 2007; Audhya *et al*, 2007), and report its function for ER-phagy during acute heat stress. We discovered that the long isoform RET-1d interacts with the LGG-1/GABARAP protein and promote the biogenesis of autophagosomes. RET-1d is mainly expressed in the nervous system and the epidermis and its depletion results in a delay in the autophagic response and a decrease in the capacity of animals to adapt to heat stress.

## Results

### The reticulon protein RET-1 interacts with LGG-1/GABARAP but not LGG-2/LC3

*C. elegans* has two homologs of the ubiquitin-like autophagy protein Atg8, LGG-1 and LGG-2, with different function during autophagosome biogenesis and maturation (Manil-Ségalen *et al*, 2014; Alberti *et al*, 2010; Chen *et al*, 2017). We performed interactome analyses of LGG-1 and LGG-2 using affinity purifications followed by mass spectrometry analysis (Yi *et al*, 2016) and identified RET-1 as a putative interactor of LGG-1/GABARAP but not LGG-2/LC3 (Figure 1A). Independently, we isolated RET-1 in a yeast two-hybrid (Y2H) screen, using LGG-1 as a bait but not with LGG-2. The sixty cDNA clones picked in the Y2H screen defined a LGG-1 interaction domain (LGG-1-ID) covering 125 amino acids located upstream of the RHD, in the non-conserved region of RET-1 (Figure 1B and Supplementary Figure S1). To check the specificity of the interaction, we performed a one by one Y2H assay using either LGG-1 or LGG-2 as baits and the RET-1[LGG-1-ID] as a prey (Figure 1C). The assay confirmed that RET-1[LGG-1-ID] specifically interacts with LGG-1, but not LGG-2 and its analysis with the iLIR software (Kalvari *et al*, 2014) revealed the presence of two putative canonical LIRs (SG**F**EK**V** and DG**F**VF**I**E). The functionality of each LIR, was assessed by Y2H one by one assay after introducing specific mutations in the RET-1[LGG-1-ID] (Figure 1B and Supplementary Figure S1). Each single LIR mutant was capable of interacting with LGG-1 but the interaction was stronger when both LIRS were present and no interaction was detected when both LIR were mutated.

**Figure 1.**
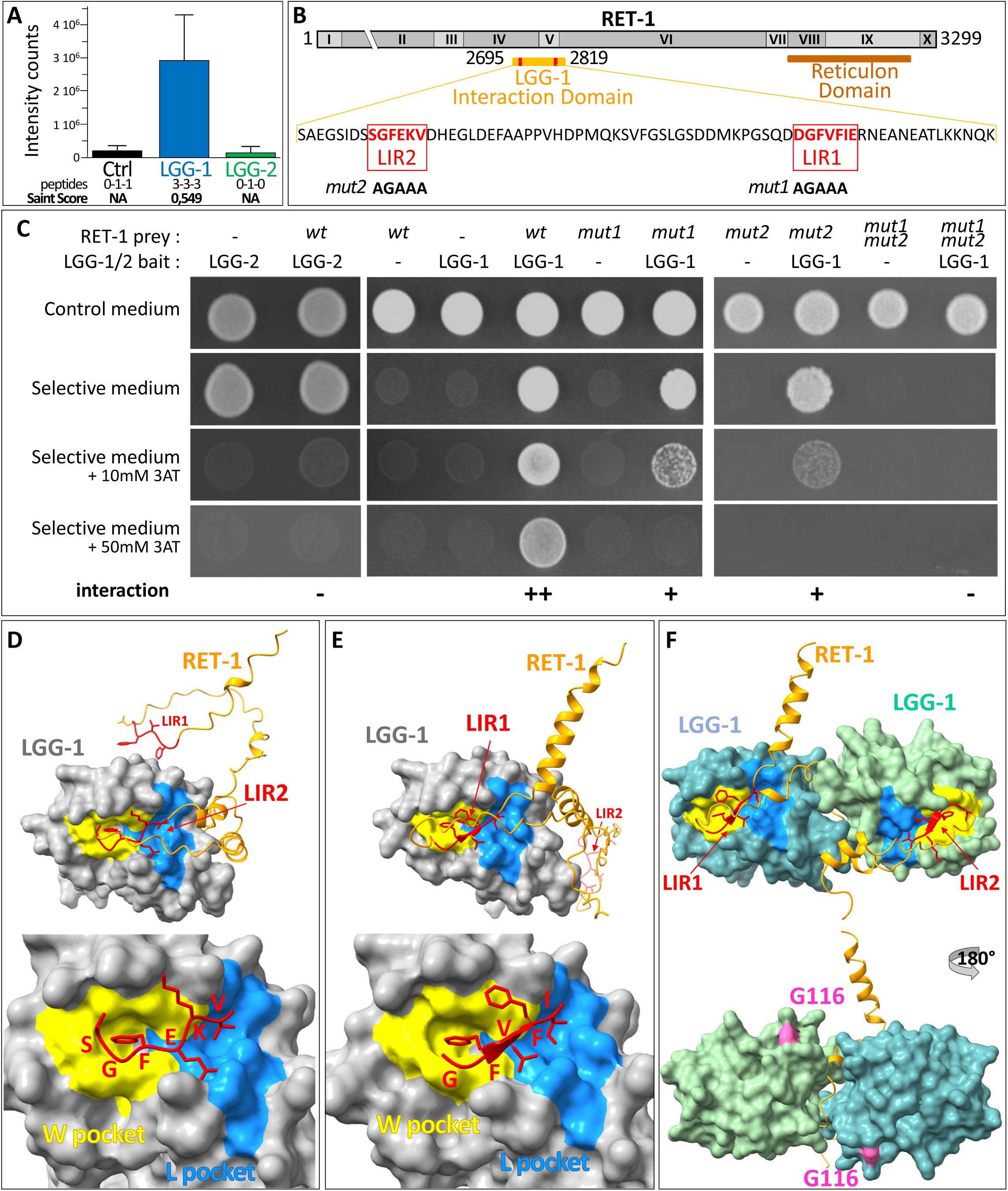
The reticulon protein RET-1 interacts with LGG-1/GABARAP but not LGG-2/LC3. (A) Mass spectrometry analysis identifies RET-1 in the interactome of LGG-1 but not LGG-2 (data from Yi *et al*, 2016). The number of peptides detected in each triplicate and the saint scores are indicated (NA non adapted). (B) Schematic representation of the *ret-1* locus according to worm base (WS269; https://wormbase.org/), with roman numbers indicating the coding exons. The position of the LGG-1 interaction domain [LGG-1-ID] of RET-1 and the reticulon domain are shown in yellow and brown, respectively. The protein sequence corresponds to the RET-1[LGG-1-ID] with the two putative LIR domains indicated in red, with the mutated sequence used for the yeast two hybrid in black bold. (C) Yeast two hybrid assays showing that LGG-1, but not LGG-2, interacts with the RET-1[LGG-1-ID] through both LIR1 and LIR2. 3-aminotriazole (3AT) is an inhibitor used to assess the strength of the interactions of the LIR domains and blocks the auto-activation of LGG-2. (D-F) AlphaFold2 modelization of the interactions of LGG-1 (grey surface) with the RET-1[LGG-1-ID] (orange backbone). Both LIR2 (D) and LIR1 (E) are predicted to interact individually to the hydrophobic pockets (yellow and blue) of LGG-1 with ipTM-score >0.84, while the global ipTM-score is 0.51 for two LGG-1 (F). G116 indicates the C terminal residue of LGG-1 I (cleaved form) involved in conjugation to phosphatidyl-ethanolamine. *See also supplementary Figure S1*.

The protein structure prediction tool AlphaFold2 (AF2) (Mirdita *et al*, 2022) was then used to simulate the interaction between LGG-1, which crystal structure has been obtained (Wu *et al*, 2015), and the RET-1 LIRs (Olsvik & Johansen, 2023). The AF2-predicted model for RET-1[LGG-1-ID] is low, suggesting an intrinsically disordered protein region. However, both LIR1 and LIR2 were predicted to interact with a high confidence score (ipTM-score 0.84 and 0.870) with the W and L hydrophobic pockets of LGG-1 (Figure 1D, E and supplementary Figure S1). The simultaneous binding of two LGG-1 molecules to RET-1[LGG-1-ID], was also simulated by AF2, but with a weaker confidence score (ipTM-score 0.51) (Figure 1F). These data indicated that RET-1 could specifically interact with LGG-1 through two LIR domains.

### LC3 Interacting Region domains are encoded in long minority RET-1 isoforms

Among the 20 independent cDNA clones identified in the initial Y2H screen, one was missing the LIR1, suggesting that alternative splicing events could generate RET-1 isoforms with 2, 1 or no LIR (Supplementary Figure S1). According to Wormbase (WS269; https://wormbase.org/), the *ret-1* locus is composed of 13 exons expended over 18 kb, which from alternative splicing and differential promoters encode at least 8 predicted isoforms that range from 204 to 3303 amino acids. To explore the relative quantification of *ret-1* isoforms, we first performed a bioinformatics analysis of a compendium of RNA-seq data for quantitative measurements of alternative splicing (Tourasse *et al*, 2017) (Figure 2A). The analysis revealed multiple isoforms forming two main categories of RET-1 proteins, the majority short forms mainly compose of the RHD, and the minority long forms. The RET-1[LGG-1-ID] is located in long isoforms with LIR1 coded by the alternatively spliced exon 5.

**Figure 2.**
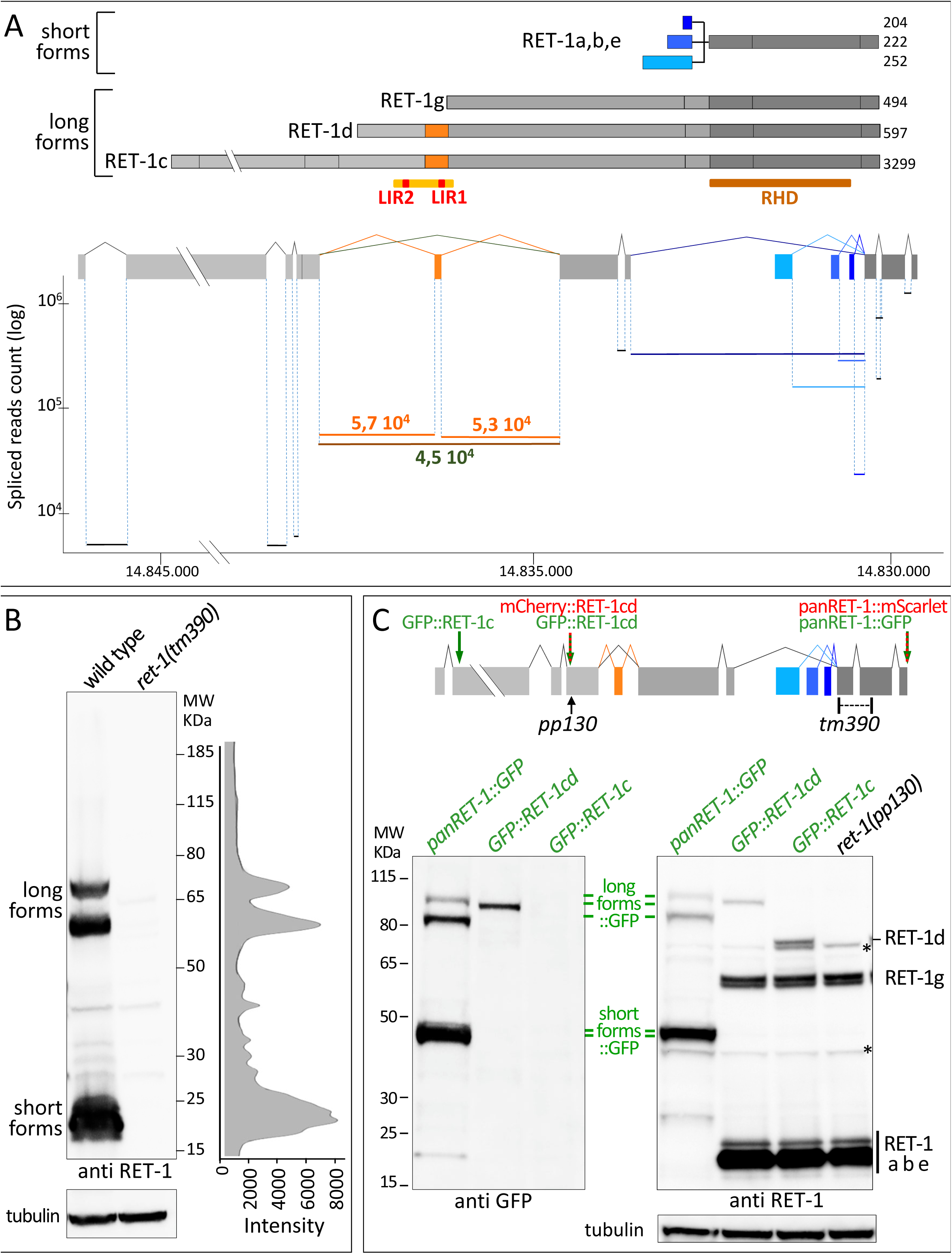
The LIR domains of RET-1 are present in the minority long isoforms. (A) The *ret-1* locus codes for short and long isoforms of RET-1 (upper). Exons are shown in grey excepted for the alternative spliced exons that are colored in blue or orange. RHD brown bar represents the reticulon homology domain and the yellow and red bar represents the RET-1[LGG-1-ID] and LIR1 and LIR2 domains. Only the main isoforms are indicated here. The quantitative analyses of the splicing events (lower part) indicate that long isoforms are minority and half of them contains the exon 5 (bold orange numbers) compared to the spliced isoform (black bold number). (B) Western blot analysis of the RET-1 isoforms expression in the wild type or the null mutant *ret-1(tm390)*. Signal quantification of each category of isoforms from the wild type extract indicates that short isoforms are majority (histogram) and RET-1c is not detected. (C) Schematic representation of the fluorescent fusions (green, red arrows) and the mutants used in the study (upper). The Western blot analysis of the RET-1 isoforms validates the expression of the GFP fusion strains and the specificity of *ret-1(pp130)* allele to deplete long isoforms (lower). The same blot was incubated successively with anti GFP antibody and anti RET-1 antibody directed against the RHD domain, and anti tubulin was used as a loading control. Asterisk indicates non specific bands.

The expression of the various isoforms by western blot analysis confirmed that the short forms were the major isoforms and that the long isoforms containing the RET-1[LGG-1-ID] represented less than 15%. Among them more than 90% corresponded to the isoforms d (thereafter called RET-1d), half of them presenting the two LIRs and half of them only one due to exon 5 skipping (Figure 2A, B). The very long isoform RET-1c, which was predicted to be very rare by the splicing analysis, was not detected by western blot. These data support RET-1d as the main candidate for an ER-phagy receptor in *C. elegans*.

### The long isoform RET-1d is expressed in the nervous system and epidermis

To confirm the *ret-1* locus analyses, fluorescent proteins tagging of long RET-1 isoforms or of whole isoforms (panRET-1) were generated using CRISPR-Cas9, as well as specific mutants (Figure 2C). The expression of GFP::RET-1cd, GFP::RET-1c and panRET-1::GFP was analyzed by western blot, which validated GFP::RET-1d and panRET-1::GFP but failed to detect GFP::RET-1c (Figure 2C).

The tissue specificity of RET-1d was deduced by the analyzes of the expression patterns of GFP::RET-1cd with GFP::RET-1c, and compared with panRET-1::GFP (Figure 3). In adult animals, panRET-1 was ubiquitously expressed and localized to a dense network, reminiscent of the ER, whose shape and intensity was variable between tissues (Figure 3A, D, E) (Lee, 2021). In contrast, GFP::RET-1cd was enriched in the nervous tissues and particularly in the neurites decorating the nerve ring and the cords (Figure 3 B, F, G). It was also expressed in the epidermis, the intestine and the body wall muscles. The longest isoform GFP::RET-1c was only detected at a very low level in the body wall muscles (Figure 3 C, H, I) raising the possibility that it is specific of the sarcoplasmic reticulum. The analyses of embryo and larvae confirmed the tissue specificity of RET-1c and d, and the colocalization of RET-1 with the ER luminal marker SP12::mCherry, indicating that the GFP tagged proteins localized to the ER. These data revealed that RET-1d is the only isoform containing the [LGG-1-ID] in neurons and epidermis and for the remainder of this study, we focussed on both tissues.

**Figure 3.**
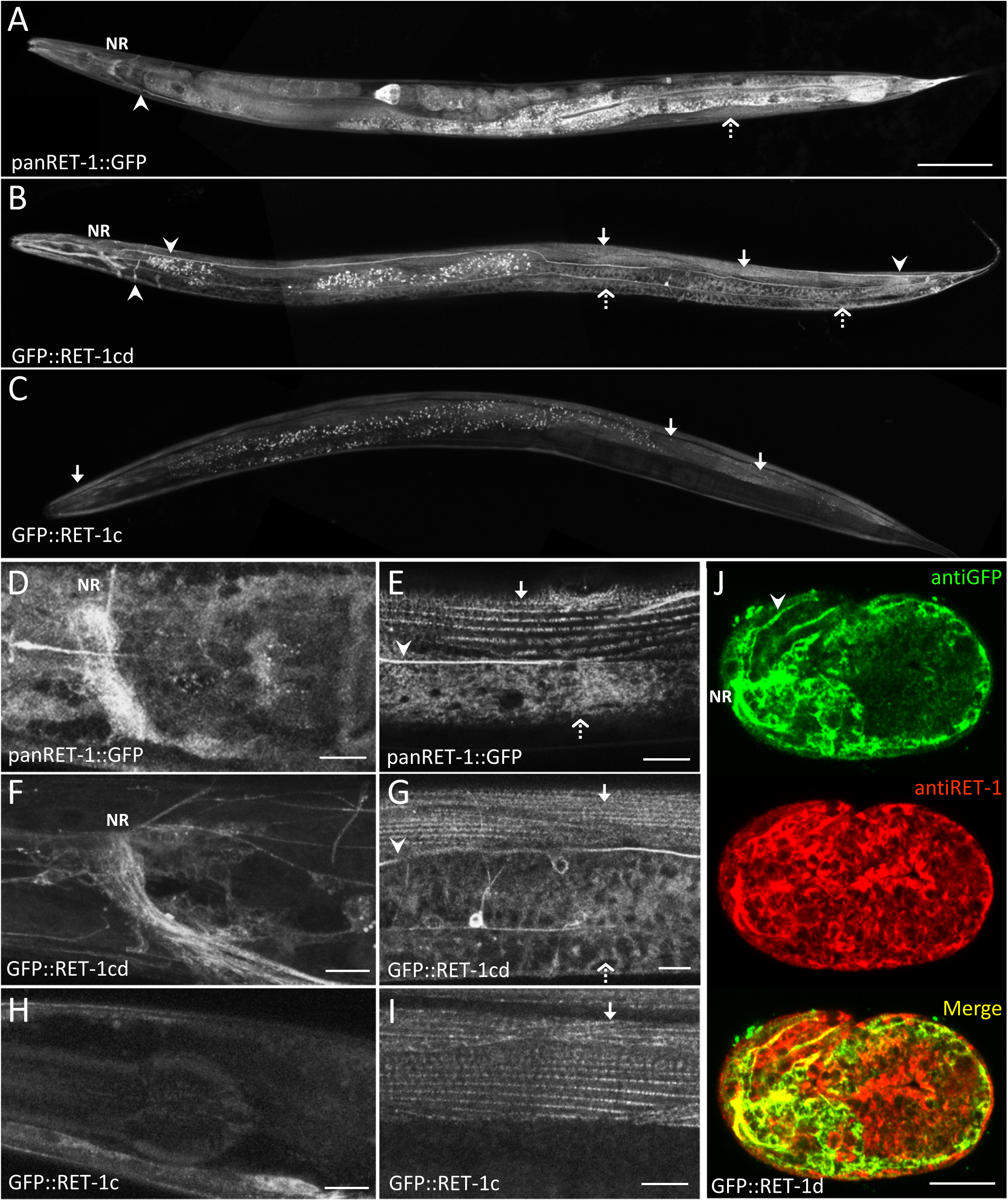
RET-1d is enriched in the nervous system and epidermis. (A-C) Assemblage of confocal images projections of live adult hermaphrodites. panRET-1::GFP (A) is ubiquitously expressed forming a network whose shape and intensity vary between tissues, RET-1cd (B) is enriched in neurons (arrowheads and nerve ring, NR) and also detected in the intestine, the epidermis (small arrows) and muscles (dotted arrows), while expression of GFP::RET-1c (C) is very weak and restricted to muscles. (D-I) Maximum projection of confocal images of the head and the body showing the dense network pattern of panRET-1 and RET-1d. (J) Confocal images of a GFP::RET-1d embryo, immunostained with antiGFP (green) and antiRET-1 (red) antibodies. The RET-1 antibody is directed against the C-terminus of the protein, present in all isoforms. While panRET-1 signal is ubiquitous, RET-1d is mostly detected in the neurons and the epidermis. *See also supplementary Figure S2*. Scale bar is 100µm (A-C) and 10 µm (D-J).

### RET-1d colocalizes with LGG-1 during ER-phagy

Next we investigated the relationships between RET-1 and LGG-1 upon induction of autophagy. We have previously shown that after acute heat stress (aHS, 60 minutes at 37°C) the ER was affected and a massive autophagy flux was triggered in L4 larvae or adults (Chen *et al*, 2021). One hour after aHS the localization of GFP::RET-1d and panRET-1::GFP in the axonal commissures was discontinuous suggesting that the long tubular ER characteristic of the neurites was fragmented (Figure 4 A, B). In the epidermis, the GFP::RET-1d pattern was also strongly modified forming dotted structures instead of a dense network, but panRET-1::GFP pattern was less affected suggesting a specific localization of RET-1d to particular ER domains upon aHS (Supplementary Figure S3).

**Figure 4.**
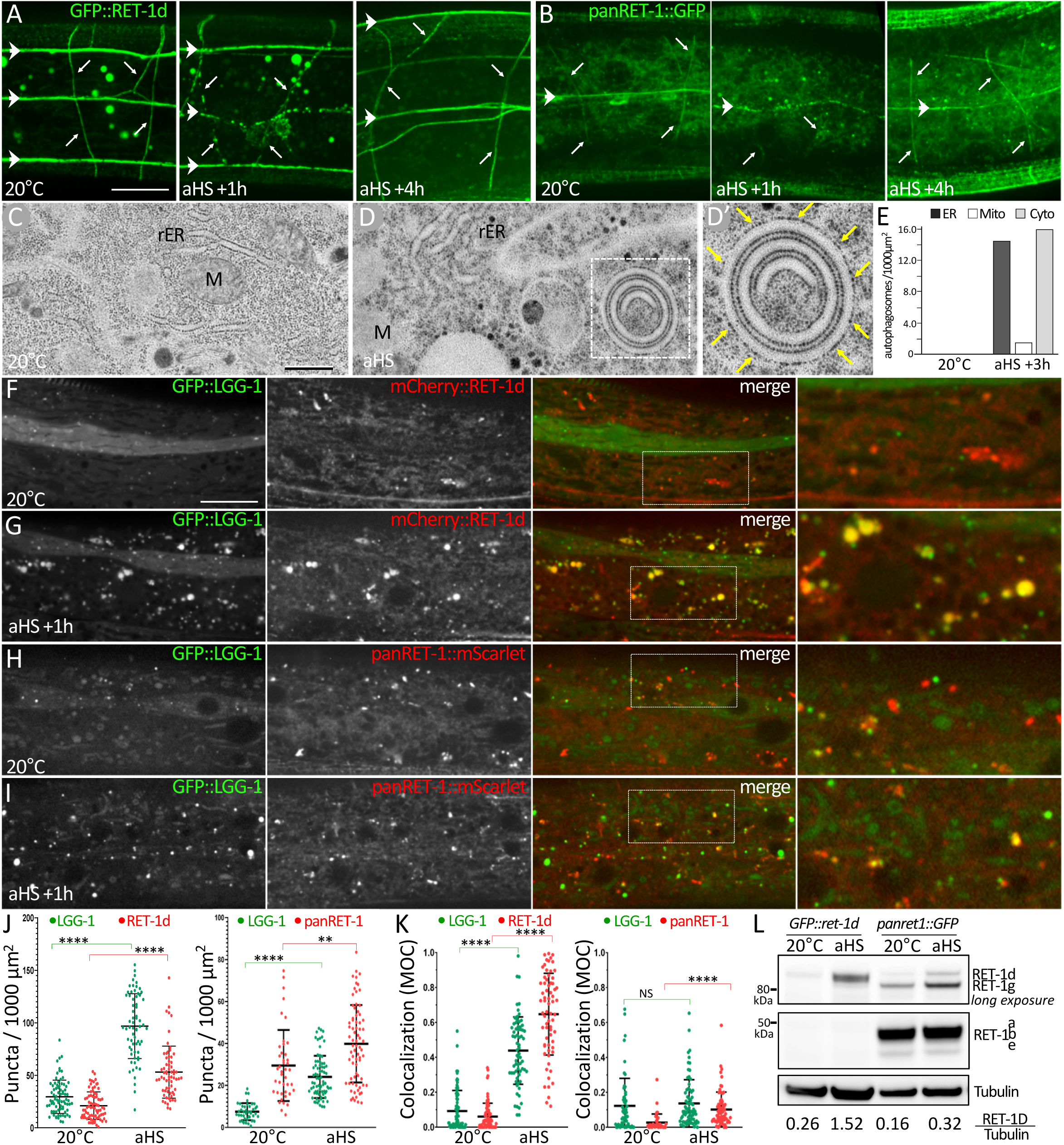
RET-1d colocalizes with LGG-1 during ERphagy. (A-B) Maximum projection of confocal images of GFP::RET-1d (A) and panRET-1::GFP (B) in the nerve cords (horizontal arrowheads) and commissures (small arrows) at 20°C or 1-4h recovery after a non lethal acute heat stress (aHS, 37°C during 60 min). The tubular ER is fragmented upon aHS and reformed after 4 hours recovery. Scale bar is 10 µm. *See also supplementary Figure S3* (C-E) Electron microscopy images of the epidermis of L4 larvae at 20°C (C) and after aHS (D, D’). The ER is mainly composed of rough ER (rER), forming tubular structures decorated by ribosomes, M indicates mitochondria. One to three hours after aHS, autophagosomes containing rER pieces are observed. Ribosomes could also be detected at the surface of the autophagosome (yellow arrows) but less regularly. Quantification of the autophagosomes containing endoplasmic reticulum (ER), mitochondria (Mito) or undefined cytoplasm (Cyto) (17 sections for 20°C, 29 for aHS) (E). *See also supplementary Figure S4* (F-I) Single confocal images of GFP::LGG-1 and mCherry::RET-1d (F, G) or panRET-1::mScarlet (H,I) in the epidermis of animal at 20°C (F, H) or one hour after aHS (G, I). (J-K) The number of RET-1d and panRET-1 puncta increases after aHS as well as LGG-1 autophagosomes (J). The Manders Overlap Coefficients between green and red pixels or red and green pixels indicate a strong colocalization after aHS between GFP::LGG-1 and RET-1d but not with panRET-1 (K). (Mean +/-SD, Mann-Whitney test, p-value****<0.0001, **<0.01, NS: non significant; n sections =78, 66, 41, 74; n animals = 36, 34, 24, 43). (L) Western blot analysis of total extracts from GFP::RET-1d and panRET-1::GFP animals with anti-RET-1 (upper panels) and anti Tubulin (lower panel). Scale bar is 10 µm (A-B, F-I) or 500nm (C, D).

Electron microscopy (EM) was then used to characterize the aspect and content of autophagosomes in the epidermis after aHS. The ER was mainly rough in the epidermis forming tubules regularly covered by ribosomes (Figure 4 C). One to three hours after aHS, inflated rER was visible in the cytoplasm, but also autophagosomes containing various cytoplasmic cargoes (Figure 4D, E Supplementary figure S4). Pieces of rER were present in 44% of the autophagosomes vesicles with variable shape and size. Inflated rER cargoes could adopt a round shape but densely packed multi-lamellar structures were also frequently observed (Supplementary figure S4). These autophagosomes often harbored ribosomes on the internal membrane and also at their external surface (Figure 4D) but generally with a lower and variable density. These observations indicated that aHS induced ERphagy in the epidermis and suggested that autophagosomal membranes could at least partially derived from rER.

To characterize the implication of RET-1d isoform in basal autophagy and induced ERphagy, the colocalization between GFP::LGG-1 and either mCherry::RET-1d or panRET-1::mScarlet was analyzed at 20°C and after aHS (Figure 4F-K). At 20°C, the number of LGG-1 positive autophagosomes was low and presented almost no colocalization with RET-1d or panRET-1 (0.03 to 0.12 Manders coefficients, Figure 4F, H, J, K). However, after aHS the number of LGG-1 puncta increased strongly as well as the colocalization with the dotted RET-1d pattern (0.44 and 0.65 Manders coefficients, Figure 4G, J, K). In contrast, the colocalization between LGG-1 and panRET-1d presented a very weak increase after aHS (0.10 and 0.14 Manders coefficients, Figure 4I, J, K). Western blot analysis of GFP:RET-1d and panRET-1::GFP also suggested a specific increase of RET-1d isoform after aHS (Figure 4 L).

Altogether, these data demonstrated that RET-1d and LGG-1 could specifically interact *in vivo* during heat stress-induced ERphagy.

### Autophagic flux is delayed in *ret-1d* mutant

To decipher the specific function of RET-1d in autophagy, we generated a mutant *ret-1d(pp130)* that depleted the RET-1 long isoforms and compared its phenotype with the mutant *ret-1(tm390)*, which affected all isoforms (Figure 2C). WB and immunofluorescence confirmed that *ret-1(tm390)* is a null mutant and that in *ret-1d(pp130)* animals only RET-1d but not the short isoforms of RET-1 was absent (Figure 2 and Supplementary figure S5). Autophagy was first analyzed in the nervous system using a GFP::LGG-1 expressed under the *rgef-1* promoter (Gelino *et al*, 2016). Wild-type, *ret-1(tm390)* and *ret-1d(pp130)* animals were submitted to aHS and GFP::LGG-1 puncta were quantified in the head neurons (Figure 5A-G) and compared with unstressed animals. In basal conditions, the pattern of LGG-1 puncta was very similar between both *ret-1* mutants and the control and quantification revealed a very weak increase in *ret-1d(pp130)* (106.0 vs 101.7) (Figure 5A-C, G). One hour after aHS, control animals presented a 22.4% increase in LGG-1 puncta, which indicated that aHS triggered autophagy in the neurons similarly to what have been described in another heat stress model (Kumsta *et al*, 2017). Surprisingly, in both *ret-1(tm390)* and *ret-1d(pp130)* the number of LGG-1 puncta was lower than in the control after aHS (37.4% and 32.2%, respectively) and the quantification revealed a decrease of 20.3% compare with the non stressed animals (Figure 5D-G). This result indicated that the autophagic response was affected in neurons when RET-1d was depleted and suggested a defect in the induction. The autophagy flux was further studied 1 to 3 hours after aHS in the epidermis, where LGG-1 autophagosome formation is massive and rapid (Chen *et al*, 2021). In wild-type animals, the number of LGG-1 puncta was maximum 1-2 hours after the aHS (> 7-fold increase) and close to the basal level at 3 hours (Figure 5 H, K). At 20°C, *ret-1(tm390)* and *ret-1d(pp130)* mutants did not show a significant difference in basal autophagy with the control. However, both mutants presented a delay in the induction and a significantly weaker number of LGG-1 puncta (< 5-fold increase) (Figure 5 I-K) without any obvious defect in their appearance.

**Figure 5.**
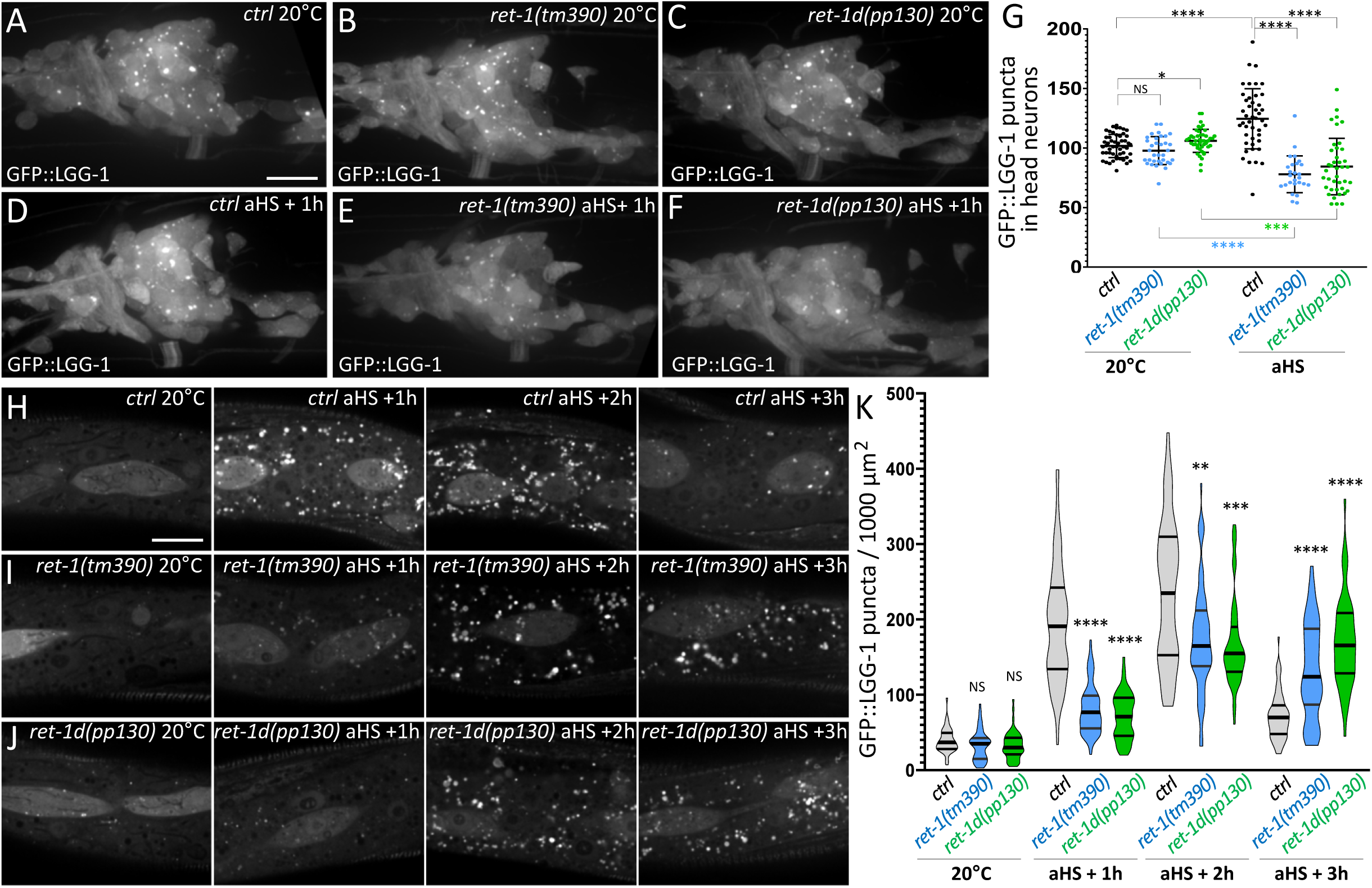
The autophagic flux is delayed in *ret-1d* mutant. (A-F) Projections of confocal images of *rgef-1p::*GFP::LGG-1 in the head neurons of control (A, D) *ret-1* null (B, E) and *ret-1d* (C, F) animals at 20°C or after aHS. (G) The quantification of puncta shows a decrease in the number of autophagosomes in both *ret-1* mutants. (Mean +/-SD, Student t test with Welch correction p-value ****<0.0001, ***<0.001, *<0.05, NS: non significant; n= 51, 34, 45, 44, 26, 48 from at least 3 independent experiments). (H-J) Single confocal images of *lgg-1p*::GFP::LGG-1 in the epidermis of control (H) *ret-1* null (I) and *ret-1d* (J) animals at 20°C or 1-3h recovery after aHS. (K) Violin representation of the number autophagosomes showing that the induction of autophagosomes is delayed in *ret-1* mutants. Bars indicate the median, first and third quartiles, n=, Kruskal Wallis test p-value****<0.0001, ***<0.001, **<0.01, NS: non significant; n sections =46, 58, 55, 72, 81, 49, 60, 68, 53, 71, 45, 50; 1 or 2 sections were measured on > 25 animals from at least 2 independent experiments). *See also supplementary Figure S5* Scale bar is 10 µm.

These analyses indicated that in absence of RET-1d the autophagy flux was diminished and suggested that RET-1d is involved in the induction of autophagosome biogenesis. Altogether, these data support a specific role for RET-1d during heat stress induced ERphagy as an inducer of autophagosome biogenesis.

### Adaptation to heat stress is defective in *ret-1d* and *lgg-1* mutants

We have previously shown that the autophagic flux induced by aHS had a protective effect on the worms, enabling the developmental recovery and participating to adaptation (Chen *et al*, 2021). The adaptation capacity of *ret-1d(pp130)* mutant was compared with *lgg-1(pp141)* mutant and wild-type animals. The *lgg-1(pp141)* mutation G116AG117STOP blocks the autophagic but not the developmental function of LGG-1 (Leboutet *et al*, 2023). In absence of stress, after 24 hours, all L4 larvae had reached the fertile adult stage and no size difference was detected between *ret-1d* or *lgg-1* mutants and wild type animals (Figure 6A, C, E, H). However, 24 h after aHS, both *ret-1d* and *lgg-1* mutants were significantly smaller that the wild type animals, indicating that the depletion of RET-1d had a detrimental effect similar to the blockage of LGG-1 autophagic function, for recovery after aHS (Figure 6B, D, F, H).

**Figure 6.**
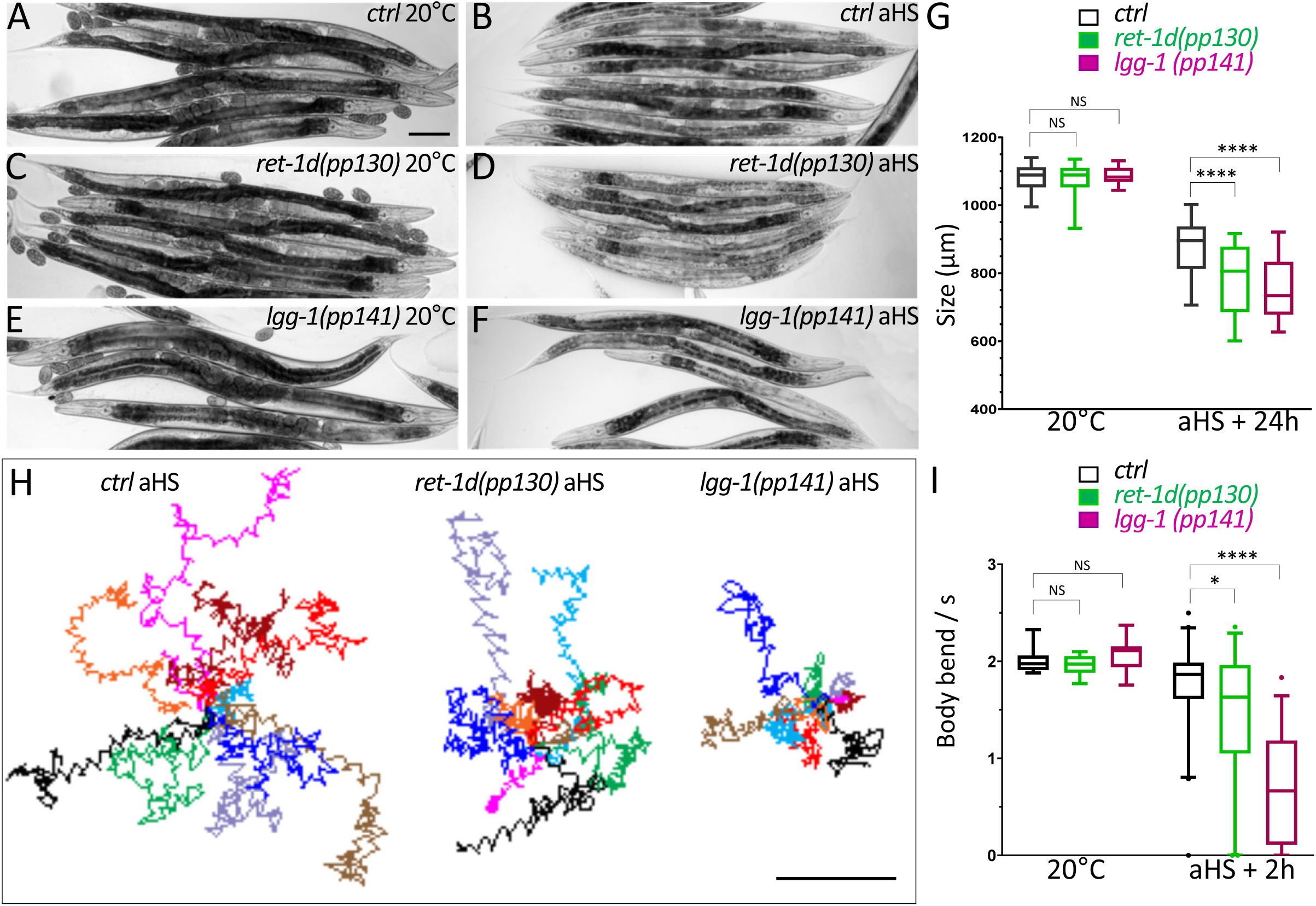
The adaptation to heat stress is impaired in *ret-1d* and *lgg-1* mutants. (A-G) The developmental delay induced by aHS is increased in *ret-1d* and *lgg-1* mutants. Early L4 larvae were maintained at 20°C (A, C, E) or submitted to a non lethal aHS (B, D, F 37°C during 60 min) and measured after 24h recovery at 20°C. (G) Boxplot of the size of the animals. One representative of three independent experiments is shown (min-max with median, first and third quartile, Mann-Whitney test p-value ****< 0.0001, NS: non significant; n = 43, 43, 25, 121, 133, 79). (H-I) The locomotion recovery after aHS is affected in *ret-1d* and *lgg-1* mutants. The motility of the worms was filmed during a 30 seconds swimming assay, 2 hours after aHS. All color lines, which represent an individual worm swimming during 10 seconds, were juxtaposed on their starting point (H). (I) Box plot of the number of body bends per second in control and heat stress conditions. One representative of three independent experiments is shown (5-95%, with median, first and third quartile, Mann-Whitney test p-value **** < 0.0001, *<0.05, NS: non significant; n = 10, 10, 12, 40, 39, 35). *See also supplementary figure S6*. The scale bar is 100 μm (A-F) or 500 µm (I).

Because RET-1d is enriched in the nervous system, we also explored the locomotion recovery after aHS. Locomotion was another hallmark for adaptation to stress because the worms were immobile after aHS and recovered their ability to move several hours later. The swimming behavior of *lgg-1(pp141)*, *ret-1d(pp130)* mutants was filmed on individual animals and the number of body bends quantified (Figure 6H, I). In absence of stress, no difference of locomotion was detected between *ret-1d* or *lgg-1* mutants and wild type animals (circa 2 bends/s, Figure 6 I and Supplementary Figure S6). Two hours after aHS, wild-type animals presented a mean swimming capacity of 1.7 bends/s with a high variability. *lgg-1(pp141)* mutants displayed a very reduced swimming capacity (0.7 bends/s) indicating that autophagy was important for locomotion recovery after aHS. Noticeably, *ret-1d(pp130)* mutants showed a significant but less pronounced decrease in swimming (1.4 bends /s), confirming a defect in recovery after aHS.

Altogether, this study demonstrated that acute heat stress induces ERphagy and a colocalization between the long reticulon isoform RET-1d and LGG-1/GABARAP. RET-1d participates to the autophagosome biogenesis induction and is involved in the adaptation to aHS.

## Discussion

ER-phagy or ERLAD (ER-to-lysosome-associated degradation) are generic terms that encompass three different modes of delivery to the lysosome: the macro-ER-phagy, the micro-ER-phagy, and the vesicular delivery pathway (Chino & Mizushima, 2023; Reggiori & Molinari, 2022). There is not yet enough studies to have an exhaustive view of the various ER-phagies but it is already clear that these processes are linked to the cargoes to be degraded and vary depending on specific ER stresses.

It is not unexpected that ER-phagy is one of the stress response pathways used by the cell to resist and adapt to heat, but this study is the first report that aHS induces a selective ER-phagy in *C. elegans*, and involves the interaction between the long reticulon isoforms RET-1d and the LGG-1/GABARAP ubiquitin-like protein. Most studies in yeast, mammals and drosophila (Mou *et al*, 2024) have described reticulon dependent ER-phagy, in specific ER stress (tunycamicyn, dithiothreitol) or global stresses (hypoxic stress, starvation) but except in plants, ER-phagies have not been studied in the context of heat stress (Bao & Bassham, 2020). Our EM analysis revealed that macro ER-phagy occurred in *C. elegans* during aHS, with sequestration of inflated ER pieces and densely packed lamellar rER in autophagosomes. It also showed that in the epidermis, the autophagosomal membrane could partially derived from the rER. We also confirmed that autophagosomes containing mitochondria are induced by aHS, but the two cargoes are almost never observed in the same vesicle, indicating that different selective autophagies occur concomitantly. The coexistence upon aHS of different autophagosomal structures with specific cargo. These data highlight the importance of heat stress induced selective autophagy in *C. elegans*, which is a poikilothermic specie, and the versatility of autophagosomes within the cell depending of the cargoes and the subcellular localization.

Several characteristics of RET-1d identified in this study, are common with SAR described during ER phagy in mammals (Chino & Mizushima, 2020; Reggiori & Molinari, 2022). RET-1d is a long isoform containing several LIRS, with a specific interaction with GABARAP/LGG-1 vs LC3/LGG-2. RET-1d LIRs are separated from the RHD by a long intrinsically disordered region allowing the interaction between phagophore and rER despite the presence of ribosomes (Chino *et al*, 2019). The localization of RET-1d in a dotted pattern after aHS suggests a role in the fragmentation of the ER, which could involve the membrane curvature function of the RHD domain, similarly to FAM134b and RTN3 overexpression (Grumati *et al*, 2017). RET-1d is involved in autophagosome initiation and colocalize rapidly after aHS with LGG-1, which has an essential function in the early steps of autophagosome biogenesis (Manil-Ségalen *et al*, 2014; Chen *et al*, 2021). The discovery of the alternative exon 5 splicing revealed RET-1d isoforms with 1 or 2 LIRs, and raised questions on their specificities. If the presence of 2 LIRS fosters the recruitment of LGG-1 it could impact the dynamic of formation and possibly the size of the autophagosomes. Because the neurons and syncytial epidermis of *C. elegans* contain long ER tubules, it is probable that RET-1d is a SAR for tubular ER. The combination of cargo recognition and autophagosome induction functions, allows RET-1d/LGG-1 to coordinate efficiently in space and time the elimination of dysfunctional ER by autophagy. All these criteria are particularly reminiscent of the reticulon RTN3 in mammals (Reggiori & Molinari, 2022; Chino & Mizushima, 2023; Gubas & Dikic, 2022) and suggest that RET-1d, despite no homology of the N terminal part, have similar functions to the long form of RTN3. *Ret-1* is the only reticulon gene in *C. elegans* (Iwahashi *et al*, 2002), but it encodes multiple isoforms which have not yet been described, and whose specific functions are unknown (Torpe *et al*, 2017). However, based on the presence of the [LGG-1-ID], no obvious function in ERphagy can be assigned out of RET-1d and the very rare isoform RET-1c.

The limited expression pattern of RET-1d suggests that other SARs implicated in ER-phagy exist in *C. elegans* and obvious candidates are the homologs of the mammalian ER proteins SEC62, atlastin, or the cytosolic CDK5RAP3 (Sun *et al*, 2021). The regulation and mechanistic action of ER-phagy receptors differ between taxa and one cannot exclude that specific receptors exist in the nematode.

Our data revealed that RET-1d dependent autophagy is involved in the systemic adaptation to acute heat stress. Both the depletion of RET-1d and its interactor LGG-1 resulted in a decrease in the locomotion and size recovery of the worms, 2 and 24 hours after aHS, respectively. The neuronal enrichment of RET-1d suggests a prominent role of neurons in the adaptation process. Quality control pathways are important for adaptation to stress and individual neurons have been identified as key sensors of ER stress in heat stress by mediating a complex signaling with other tissues (Prahlad *et al*, 2008). A strong ER-phagy flux is probably triggered because UPR and ERAD pathways are not sufficient to cope with an acute stress (60 min, 37°C) (Lee, 2021). The dynamic and intensity of the autophagy flux upon aHS vary between tissues (Chen *et al*, 2021) and it would be interesting to identify whether a subset of neurons is involved in a systemic response. In this regard, *C. elegans* represents a good paradigm for understanding the function of ER phagy at the organismal level and the connection between reticulon and neurological disorders (Ferro-Novick *et al*, 2021; Kulczyńska-Przybik *et al*, 2021; Zou *et al*, 2018; Shi *et al*, 2014).

## Acknowledgments

The authors would like to thank K. Oegema and J. Audhya, for giving us antibody against RET-1, G. Seydoux and A. Paix for technical support concerning CRISPR-Cas9 technology. We thank the National Bioresource Project for the nematode (NBRP, Japan) and the Caenorhabditis Genetic Center (CGC) which is funded by National Institutes of Health (NIH) Office of Research Infrastructure Programs (P40 OD010440) for providing several strains used in this study. The present work has benefited from Imagerie-Gif core facility supported by l’Agence Nationale de la Recherche (ANR-10-INBS-04/FranceBioImaging; ANR-11-IDEX-0003-02/ Saclay Plant Sciences). We thank the BIOI2 platform for making the ColabFold pipeline easily accessible at the I2BC. This work was supported by the Agence National de la Recherche (grant numbers ANR-12-BSV2-018 and ANR-23-CE13-0013-01) and the Association pour la Recherche contre le Cancer (grant number SFI20111203826). V.S. is recipient of fellowship from the Ligue Nationale contre le Cancer. CS is supported by Association Nationale de la Recherche et de la Technologie (CIFRE N° 2022/0047) and AP by a CNRS Ph.D. fellowship.

## Material and Methods

### *C. elegans* culture and strains

Nematode strains were grown on nematode growth media (NGM) plates and fed with *Escherichia coli* strain OP50 (Brenner, 1974). The *C. elegans* Bristol N2 strain was used as a wild-type strain. Genotypes of all the strains used in this study are listed below.

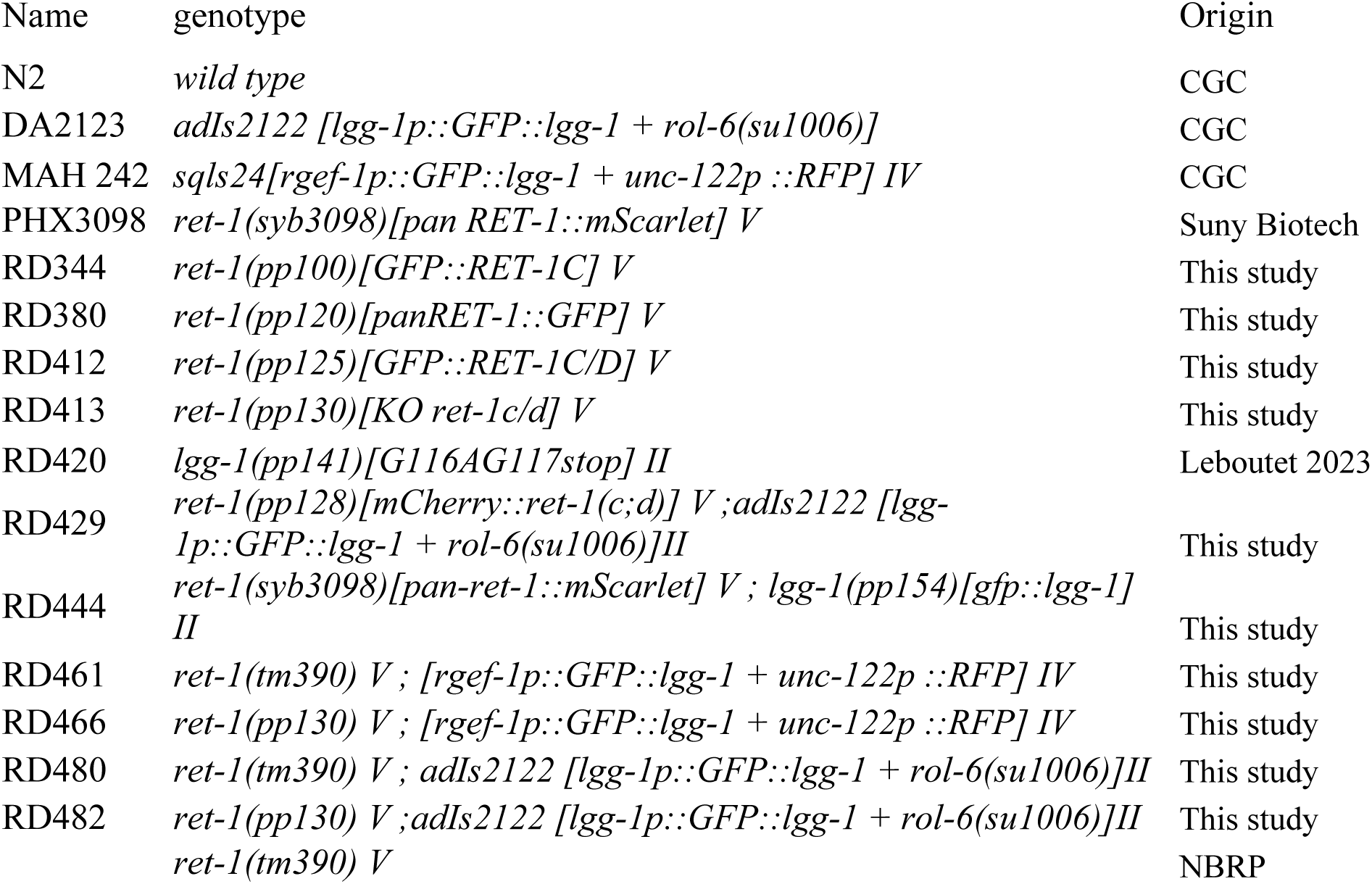

RD344, RD380, RD412 and RD413 strains were obtained by a CRISPR-CAS9 approach optimized for *C. elegans*, using a *dpy-10* co-CRISPR protocol (Paix *et al*, 2015; Leboutet *et al*, 2023). *ret-1(syb3098)* was generated by SunyBiotech (Fuzhou, China) using CRISR/Cas9 genome editing method. In RD344, GFP was inserted in-frame in exon 2 of *ret-1c*, in order to avoid secondary structures. In *pp120* or *syb3098*, GFP or mScarlet was inserted in C-terminus of *ret-1*, just before the stop codon, common to all isoforms. *pp125* correspond to an insertion of GFP in the beginning of *ret-1d* first exon. *pp130* is the result of an non homologous end joining event at the same Cas9 cutting site, inserting 125 nucleotides and a stop codon in c and d isoforms. The insertions were verified by sequencing. RD461, RD466, RD444, RD429, RD480 and RD482 have been constructed by genetic crosses.

### Heat stress

For heat stress condition, one-day-old adult worms were allowed to lay eggs for 1 hour at 20°C on NGM plates and then removed. NGM plates were maintained at 20°C until the progeny reached early 4th larval stage and were submitted to 37°C for 60 min in an incubator (Binder) followed by a recovery at 20°C for 1 to 24 hours depending on the analyses. Control worms were treated similarly but maintained at 20°C without heat stress.

For swimming analyses, worms were picked two hours after heat stress, into a drop of M9 buffer, and the thrashing was recorded for 30s (1 image per 60ms, 1.2X magnification) with a Leica MZ75 and Roper Scientific Photometrics CoolSNAPcf camera. The recordings were analyzed and quantified using the Image J plugin worm Tracker (Nussbaum-Krammer *et al*, 2015). Path tracing was based on the first 10 seconds of the movie.

The size of the worms was analyzed 24h after heat stress with a wide-field AxioImagerM2 microscope (Zeiss) and measured with Image J using segmented line from the mouth to the beginning of the tail.

### Protein extraction and Western-blot

Unsynchronized population of worms or L4 larvae after aHS+3h were harvested in M9 from OP50 plates and the pelleted worms were lysed in equal volume of PBS Triton 2% + 10% PIC (Protease inhibitor cocktail, Roche 1183617), and glass beads (425-600µm, Sigma IG8772), with Precellys 24 machine (1min 6000rpm, 5 min on ice, 1min 6000rpm, and on ice). The proteins extracts have been centrifuged at 15000rpm and separated on a NuPage 4-12% bis - tris gel (Life Technologies, NP0321BOX). Membrane Transfer has been performed with iBlot 2 machine (Thermo Fisher). The non-specific sites are then blocked after the incubation for one hour with TBS Tween 0,1% (Tris-Base, NaCL, Tween20, pH 7,5) + 3% milk. Membranes were probed with primary antibodies Anti-RET-1 (1:5000, rabbit, gift from A. Audhya), anti-GFP (1:2000, mouse, Roche 1814460) and anti-Tubulin (1:1000, mouse; Sigma, 078K4763) overnight at 4°C and revealed using HRP-conjugated antibodies (1: 5000 promega W401B and 1:5000 promega W4021) and the Super Signal Pico Chemiluminescent Substrate (Thermo Fisher Scientific, 34579). Signals were revealed on Chemidoc® Touch (Biorad) and quantified with Image Lab software.

### Immunofluorescence

Embryos were prepared for antibody staining on poly-L-lysinated (Sigma P8920) slides. The samples were freeze–fractured and fixed in desiccated methanol at -20°C for 30 minutes. After two 5min washes with PBS 1X and a saturation step with 1X PBS containing 4% BSA (Sigma A7030) and 0.1% Triton (X-100 Sigma T9284) (Abdil buffer) for 1h, samples were incubated in 1X PBS, 4% BSA and 0.1% Triton buffer overnight at 4°C with primary antibodies, in humid chamber. After four 15 min washes in 1X PBS 0.1% Triton, samples were incubated in the Abdil buffer for 2h at room temperature with secondary antibodies and a DNA labeling agent. Final mounting was carried out in DABCO (Sigma). Primary antibodies used in this study: mouse monoclonal anti-GFP at 1:250 (Roche, 1814460), chicken polyclonal anti-GFP 1:40 (Abcam Ab13970), rabbit anti-RET-1 (gift from A. Audhya) at 1:1000. As secondary conjugated antibodies, Alexa Fluor® 488, Alexa Fluor® 568 (Molecular Probes) were used at a dilution of 1:500, and Alexa Fluor® 647(anti Rabbit, Invitrogen A21244) at 1:500. DNA was labeled using Hoechst at 1/500 (Molecular Probes).

### Light microscopy

Embryos and adult worms were imaged on a confocal Leica TCS SP8 microscope with 100x and 40X objective, respectively. Live whole worm images are assemblages of projected stacks. For live imaging, larvae were mounted on a 2% agarose pad and immobilized by 40 mM sodium azide and observed in the next 30 min on a Nikon Eclipse TiE inverted microscope with Yokogawa CSU-X1-A1 spinning disc module and Prime 95B camera. Confocal images were captured with serial z sections of 0.5 μm.

Images analysis and processing was performed with imageJ (Fiji) (Schindelin *et al*, 2012)(Schindelin *et al*, 2012). The colocalization between LGG-1 and RET-1 was analyzed in the syncytial epidermis of worms excluding seam cells. The number of puncta was quantified using an object based method (center of mass with a minimum size of 3 pixels). Manders overlapping colocalization coefficients were determined with the JacoP plugin v2.1.4 (Bolte & Cordelières, 2006). The analysis of autophagy in neurons and epidermis of *ret-1* mutants was based on GFP::LGG-1 expression under the promoter *rgef-1*(MAH242) and its own promoter (DA2123), respectively. The number of the LGG-1 puncta in the head neurons was quantified with the Cell Counter plugin on a maximum intensity projection of 15 sections. For the quantification of the autophagic flux in the epidermis, the LGG-1 puncta were quantified by particle analysis after post processing by the trainable Weka Segmentation plugin (Arganda-Carreras *et al*, 2017).

### Electron Microscopy

Control or heat stressed worms were transferred to 20% BSA (Sigma-Aldrich, A7030) on 1% phosphatidylcholine (sigma-aldrich) pre-coated 200µm deep flat carriers, followed by cryo-immobilization in the EMPACT-2 HPF apparatus (Leica Microsystems; Vienna Austria) as described (Largeau & Legouis, 2019). Cryo-substitution was performed using an Automated Freeze-substitution System AFS2 (Leica Microsystems; Wetzlar, Germany). Deep flat carriers were inserted in the incubating chamber of AFS2 pre-set at -90°C with freeze substitution buffer (acetone, 1% osmium, 0,25% uranyl acetate) in cryogenics tubes. Samples were kept at -90°C for 30h then temperature sloped to -30°C with a gradient of 3°C per hour and maintained at - 30°C during 12h before going to 0°C. Sample were then removed from AFS and kept à 4°C in ice and washed 4 times 10 min in anhydrous acetone. Samples were infiltrated at room temperature with 25% for 4h, 50% for 4h, 75% over night , 100% for 8h and 100% over night of EPON (Agar Scientific, R1165) before been embedded in fresh EPON 24h at 60°C. Ultrathin sections of 80 nm were cut on an ultramicrotome (Leica Microsystems, EM UC7) and collected on a formvar and carbon-coated copper slot grid (LFG, FCF-2010-CU-50). Sections were contrasted with 2% uranyl acetate for 15 minutes and 0.08 M lead citrate (Sigma-Aldrich, 15326) for 8 minutes. Sections were observed with a Jeol 1400 TEM at 80 kV and images acquired with a Gatan 11 Mpixels SC1000 Orius CCD camera.

### Mass spectrometry

RET-1 was identified as an interactor for LGG-1, using an immunoprecipitation approach from a mixed-stage population of worms expressing GFP::LGG-1, followed by a mass spectrometry analyses. Briefly, the experiments were done in quadruplicate, and immunoprecipitates were separated into four fractions by 1D electrophoresis. Proteins were in-gel digested using trypsin, and tryptic peptides were analyzed by mass spectrometry. The data generated by the nanoLC−MS/MS analysis were further processed for protein identification and label-free quantification was performed with MaxQuant using the LFQ procedure of intensity determination of the MS signal and normalization. Data, were analyzed through the SAFER workflow (mass Spectrometry data Analysis by Filtering of Experimental Replicates). The detailed protocols of the immunoprecipitation, the protein digestion and tryptic peptide preparation as well as the nanoLC−MS/MS analysis and data processing have been reported (Yi *et al*, 2016).

### Yeast two hybrid interactions

The yeast two-hybrid interaction assays were performed by Hybrigenics Services SAS, Paris, France. RET-1 was initially identified in the ULTImate Y2H screening of LGG-1 against the *C. elegans* mixed stage cDNA. library. One-by-one interaction assays were first used to test for protein interaction between LGG-1 or LGG-2 and RET-1, then between RET-1 mutated for the LIR1 or LIR2 motif and LGG-1. Briefly, open reading frames of LGG-1 and LGG-2, were amplified by PCR and fused to LexA DNA binding domain. The prey fragment for the RET-1 was extracted from the initial screening and either the wild-type sequence or the mutated LIRs were fused with Gal4 Activation Domain. All constructs were checked by sequencing the entire insert. Bait and prey constructs were transformed in the yeast haploid cells, respectively L40 Gal4 (mata) and YHGX13 (Y187 ade2-101::loxP-kanMX-loxP, matα) strains and the diploid yeast cells were obtained using a mating protocol. Positive yeast two-hybrid interactions were based on the HIS3 reporter gene. Interaction pairs were tested in duplicate as two independent clones from each diploid were picked for the growth assay. Serial dilutions (10^-1^ to 10^-4^) of a 5.104 diploid cell culture expressing both bait and prey constructs were spotted on several selective media for growth control and interaction. 3-aminotriazole (3-AT), an inhibitor of the HIS3 gene product, was added at increasing concentrations (1 to 50 mM) to increase stringency and reduce auto-activation by LGG-2.

### Modelization

The Alphafold 2.3 modelization of the interactions were obtained using the ColabFold v1.5.2 pipeline (Mirdita *et al*, 2022) and MMseqs2 verified version, on the cluster Alphafold2I2BC. A fragment of the first 142 amino-acids of RET-1d (covering the RET-1[LGG-1-ID]) and the full length LGG-1 I form were used to generate 5 models, with 3 recycles on environmental databases in sequence mode (MGnify). Unpaired plus paired mode were used for multimer predictions.

### Splicing analysis

The quantitative exon usage information for the *ret-1* locus was obtained from the annotated gene models for the entire *C. elegans* genome described previously (Tourasse *et al*, 2017). Briefly, the relative abundance of each splicing event was determined based on a compendium of RNA-seq data and an automated curation method taking into account the expression level of each gene to discriminate robust splicing events from biological noise.

### Statistical analyses

All statistical analyses were performed using either GraphPad Prism or the R-software. The Shapiro-Wilk’s test was used to evaluate the normal distribution of the values. Data derived from different genetic backgrounds were compared by Student t test, Kruskal-Wallis or Wilcoxon-Mann-Whitney tests. Error bars are standard deviations (SD). Boxplot and violin plot representations indicate the minimum and maximum, the first (Q1/25th percentile), median (Q2/50th percentile) and the third (Q3/75th percentile) quartiles. Statistical tests used, with p-values, the number of samples are indicated in the figure legends. All experiments have been done at least three times

---

**Supplementary Figure S1-related to Figure 1.**
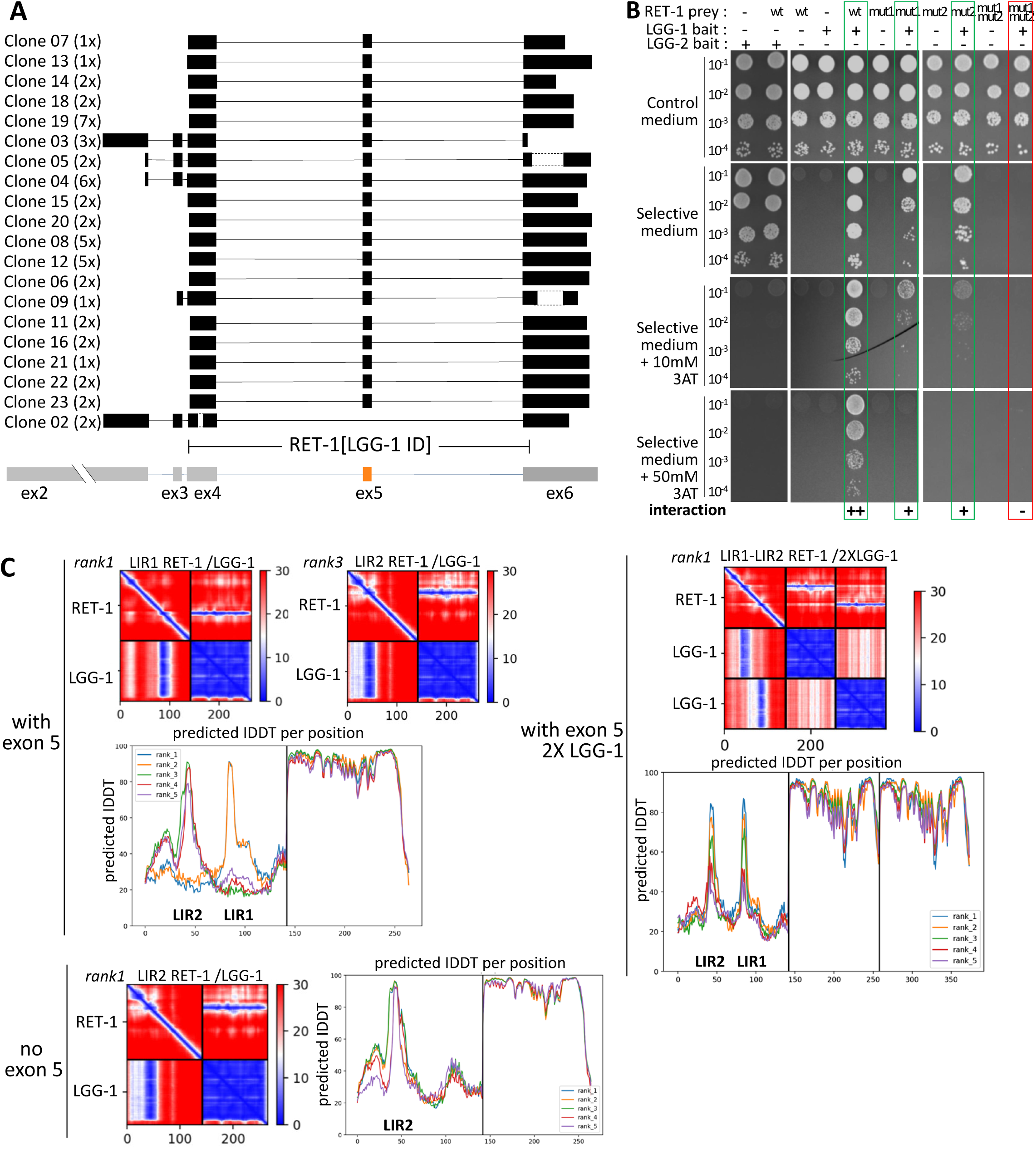
LIR1 and LIR2 domains of reticulon protein RET-1 interact with LGG-1. (A) Analyses of *ret-1* cDNA clones from the yeast two hybrid screen identified a minimal LGG-1 interaction domain [LGG-1-ID] covering exon 4 to 6, and exon 5 (orange) as a putative alternative spliced exon. (B) The one to one Y2H interaction was tested on two different clones (one is shown here) with serial dilutions in presence of 3-aminotriazole (3AT). (C) AlphaFold2 prediction aligned error (PAE) score and predicted local distance difference test (lDDT) for the models of LIR1/LGG-1; LIR2/LGG-1 and both LIR1-2/LGG-1 (2X) shown in Figure 1 D-F, In absence of exon 5, only interaction with LIR2 site is simulated.

**Supplementary Figure S2-related to Figure 3.**
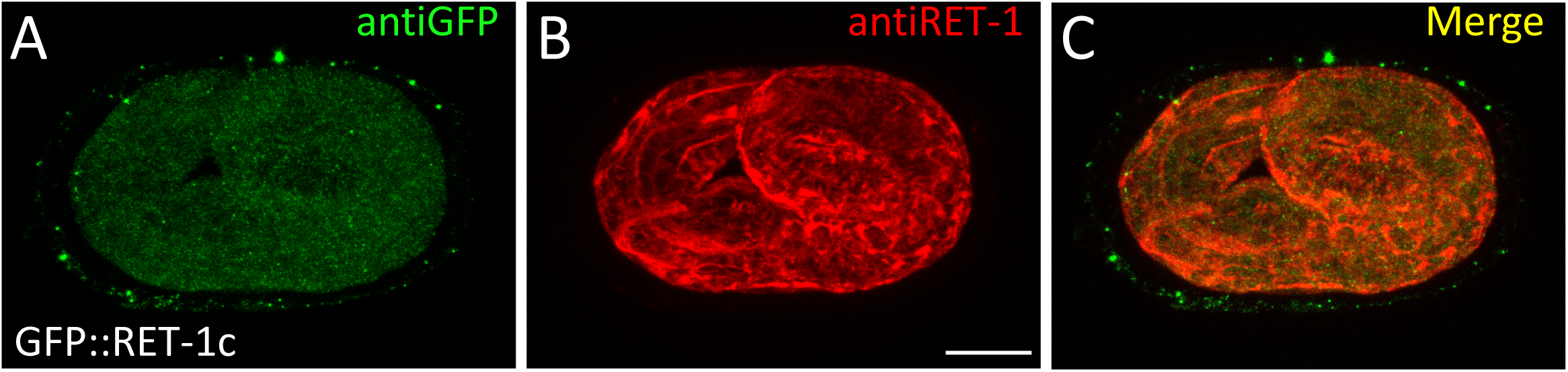
RET-1c isoform is not expressed in the embryo. Maximum projection of confocal images of a GFP::RET-1c embryo immunostained with anti GFP antibody (A) and anti RET-1 (B) antibody that recognizes all isoforms, and merge (C). Scale bar is 10µm.

**Supplementary Figure S3-related to Figure 4.**
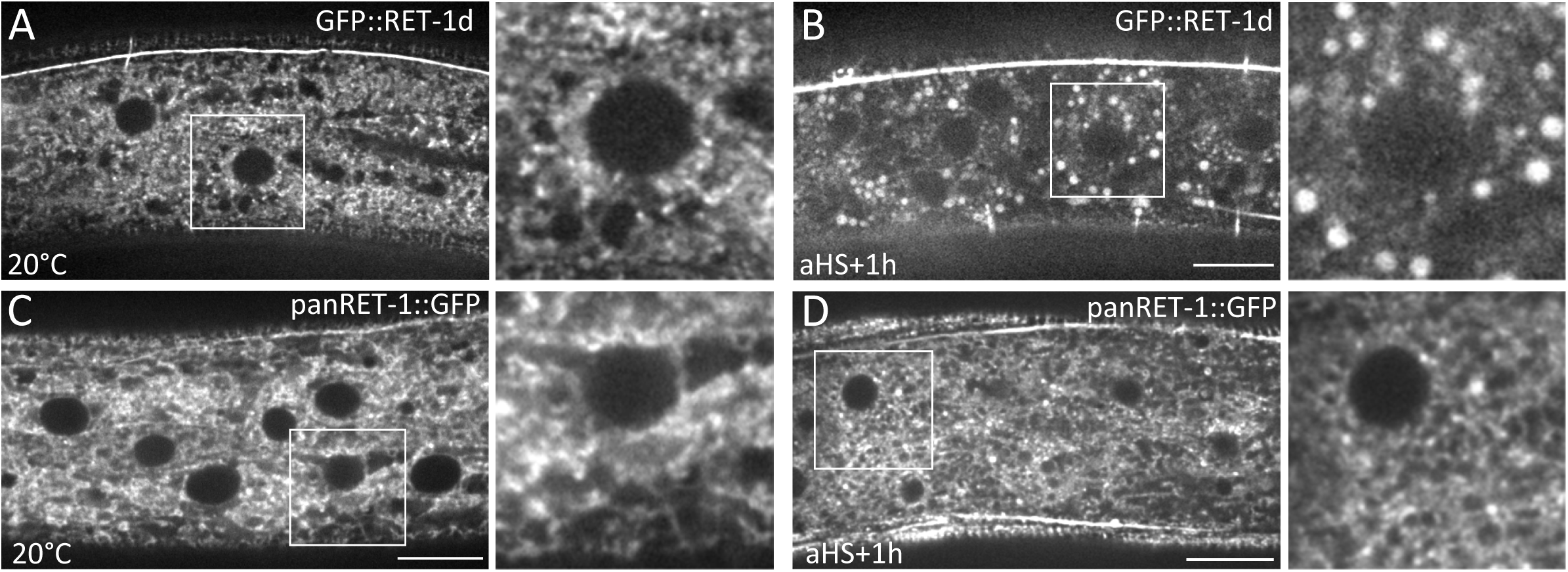
RET-1d but not panRET-1 localizes to puncta after aHS. Single confocal section of the epidermis of GFP::RET-1d (A, B) or panRET-1::GFP (C, D) at 20°C (A, C) or after aHS (B, D). The white square indicates the region around an epidermal nucleus, which is magnified on the right panel. After aHS GFP::RET-1d is mainly localized to puncta while panRET-1::GFP remains localize to a meshwork with few puncta. Scale bar is 10µm.

**Supplementary Figure S4-related to Figure 4.**
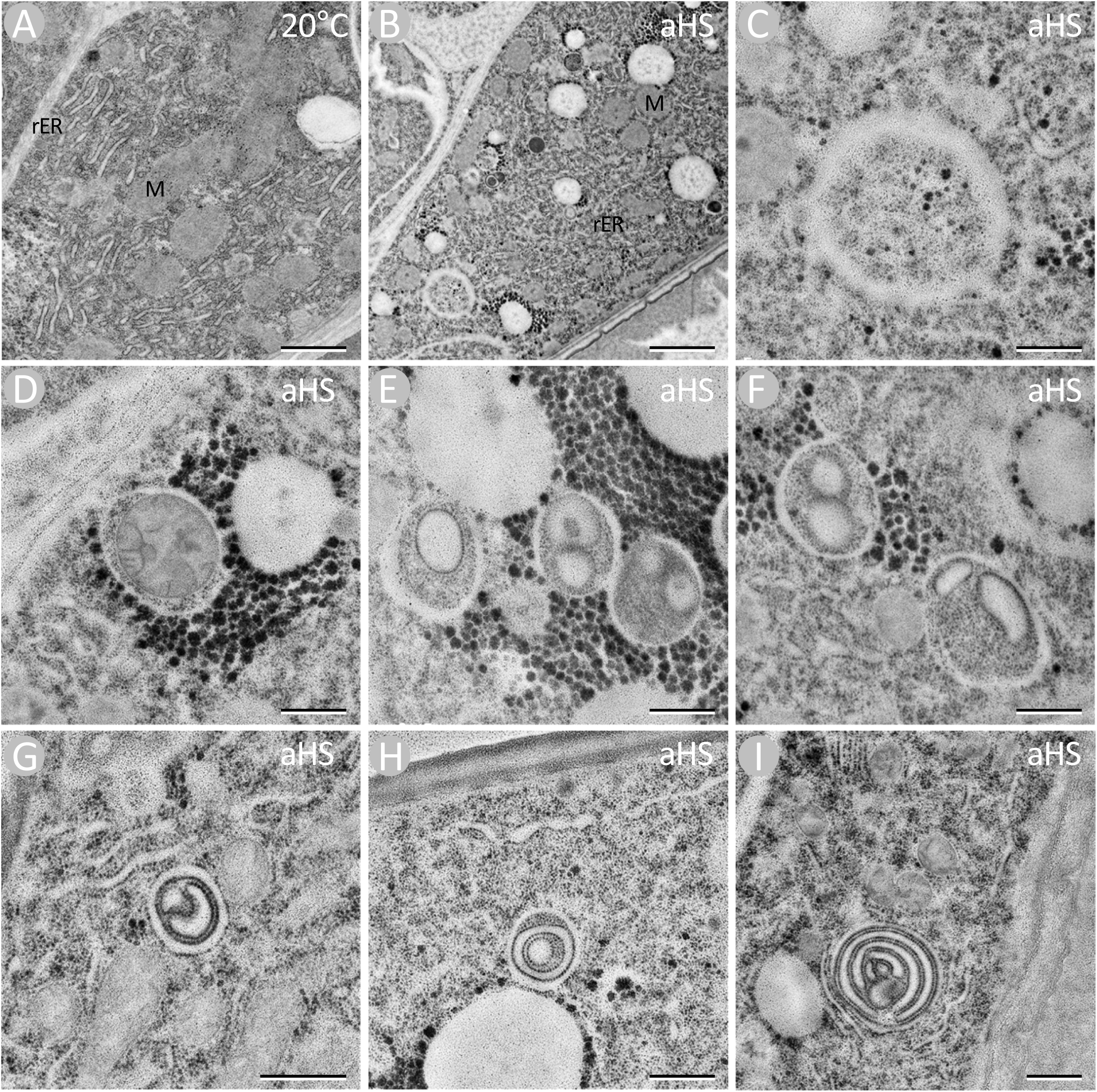
ER pieces are present in autophagomes after aHS. Electron microscopy images of the epidermis of L4 larvae at 20°C (A) and after aHS (B-I), rER indicates rough ER and M a mitochondria. (C, D) An autophagosome with undefined cytoplasmic content (C) and a stressed mitochondria with characteristic dense structures in the matrix (D). (E-I) Autophagosomes containing inflated rER pieces that can adopt a round aspect or a multi-lamellar organization. Ribosomes are also detected at the surface of the autophagosomes (G-I) with a very variable density that is generally weaker than the internal autophagosomal membrane. Scale bar is 2 µm (A, B) or 500 nm (C-I).

**Supplementary Figure S5-related to Figure 5.**
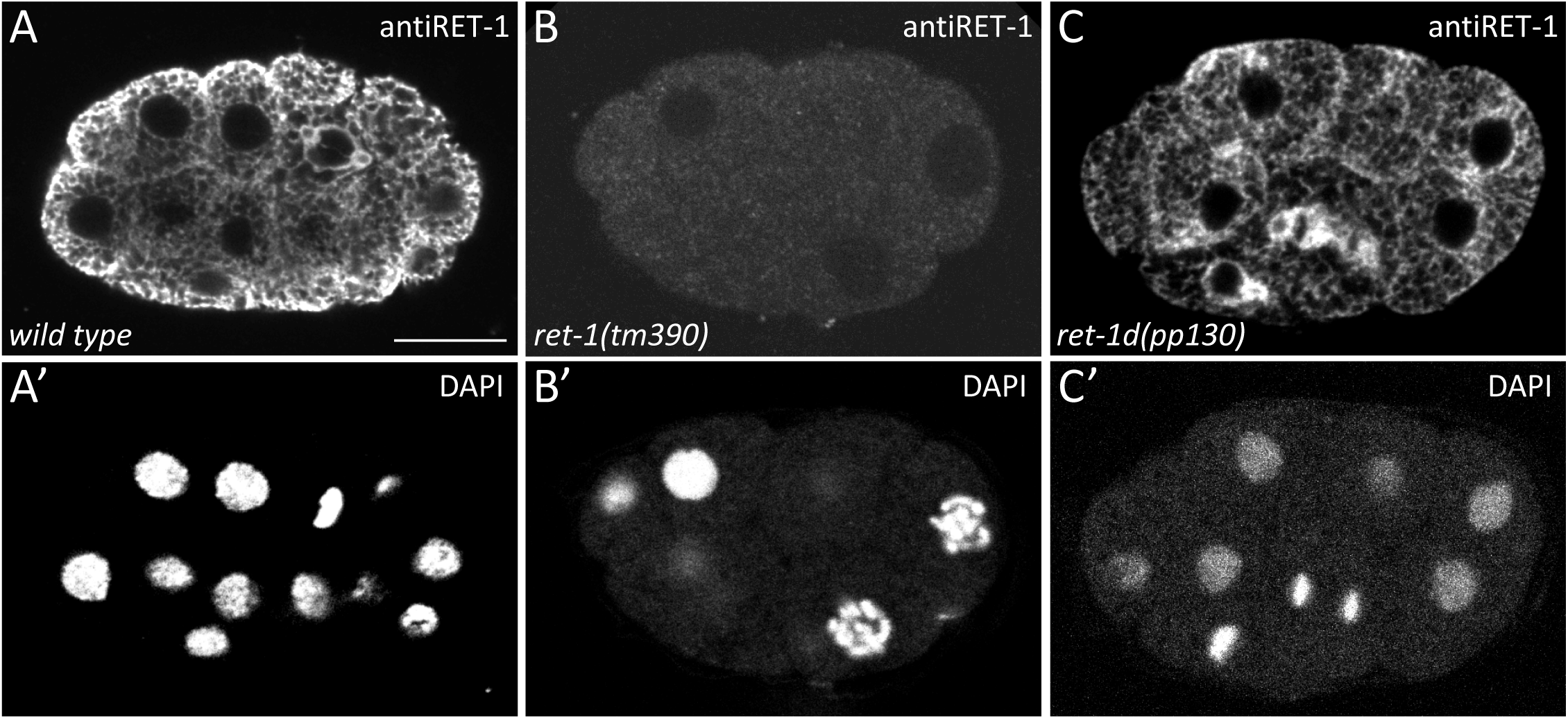
*ret-1d* mutant does not affect the localization of short isoforms of RET-1. (A-C) Single confocal image of *wild type* (A) *ret-1 null* (B) and *ret-1d* (C) embryos immunostained with antiRET-1 antibody (A-C) and Dapi (A’-C’). The RET-1 antibody is directed against the C-terminus of the protein, present in all isoforms. While the *ret-1* null allele *tm390* depletes all isoforms, the *ret-1d* mutant allele *pp130* does not affect the localization of the short and majority isoforms. The scale bar is 10 μm.

**Supplementary Figure S6-related to Figure 6.**
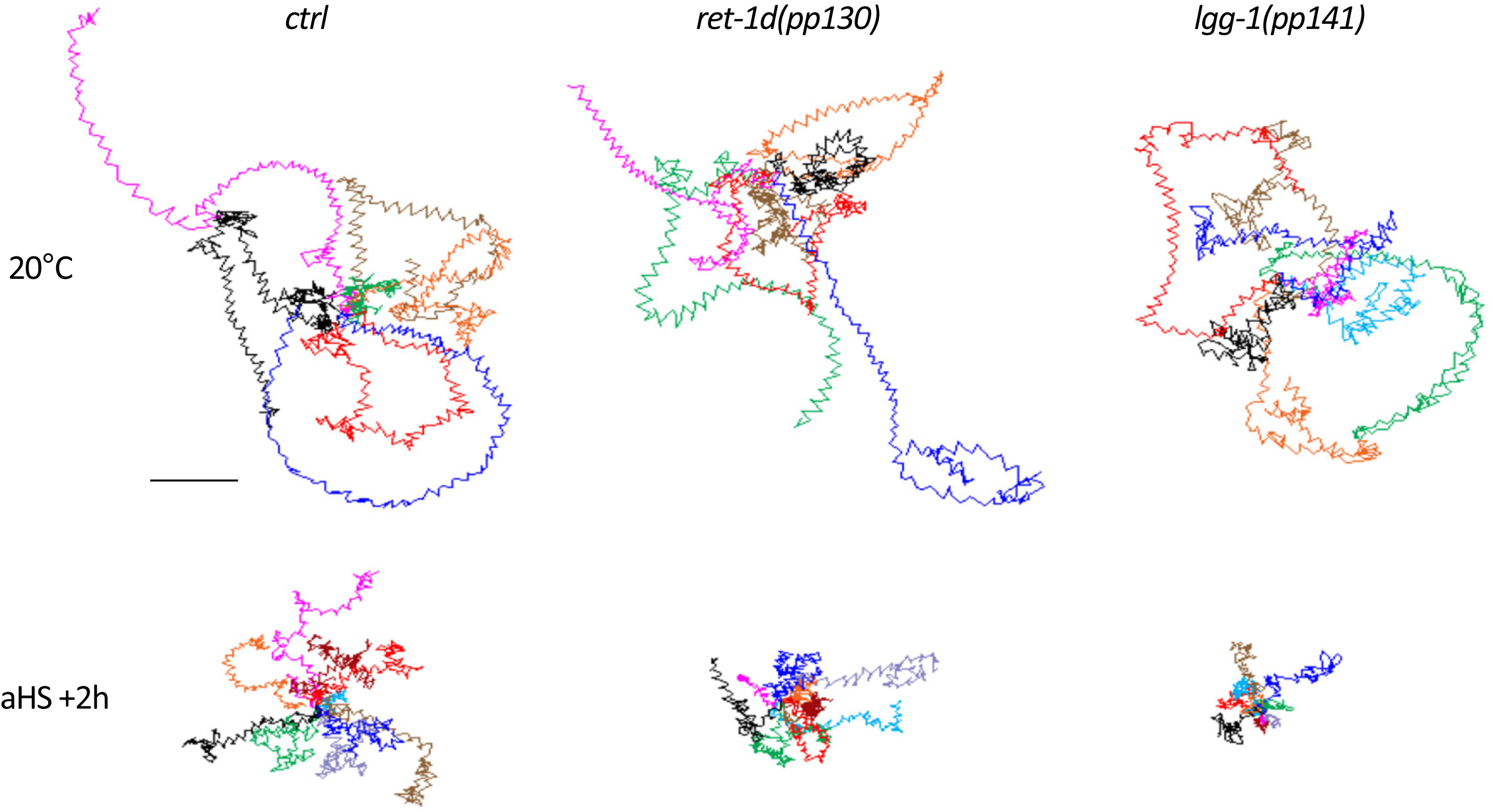
Locomotion of *ret-1d* and *lgg-1* mutants is normal at 20°C. Early L4 larvae were maintained at 20°C (upper) or submitted to a non lethal aHS (lower, 37°C during 60 min) and the motility of the worms was filmed during a 30 seconds swimming assay, 2 hours after aHS. All color lines, which represent an individual worm swimming during 10 seconds, were juxtaposed on their starting point. Images after heat stress are the same as in figure 6 H. The scale bar is 500 µm.

